# Importance of homo-dimerization of Fanconi-associated nuclease 1 in DNA flap cleavage

**DOI:** 10.1101/236208

**Authors:** Timsi Rao, Simonne Longerich, Weixing Zhao, Hideki Aihara, Patrick Sung, Yong Xiong

## Abstract

Fanconi-associated nuclease 1 (FAN1) removes interstrand DNA crosslinks (ICLs) through its DNA flap endonuclease and exonuclease activities. Crystal structures of human and bacterial FAN1 bound to a DNA flap have been solved. The *Pseudomonas aeruginosa* bacterial FAN1 and human FAN1 (hFAN1) missing a flexible loop are monomeric, while intact hFAN1 is homo-dimeric in structure. Importantly, the monomeric and dimeric forms of FAN1 exhibit very different DNA binding modes. Here, we interrogate the functional differences between monomeric and dimeric forms of FAN1 and provide an explanation for the discrepancy in oligomeric state between the two hFAN1 structures. Specifically, we show that the flexible loop in question is needed for hFAN1 dimerization. While monomeric and dimeric bacterial or human FAN1 proteins cleave a short 5’ flap strand with similar efficiency, optimal cleavage of a long 5’ flap strand is contingent upon protein dimerization. Our study therefore furnishes biochemical evidence for a role of hFAN1 homodimerization in biological processes that involve 5’ DNA Flap cleavage.

## INTRODUCTION

Interstrand DNA crosslinks (ICLs) interfere with DNA replication and transcription. Failure to remove ICLs can induce cell cycle arrest, cell death, and genome instability. The Fanconi-associated nuclease 1 (FAN1) is a DNA structure-specific nuclease involved in ICL repair [1-4]. It possesses both 5’ flap endonuclease and 5’ to 3’ exonuclease activities. In cells, FAN1 likely functions in concert with other nucleases to unhook ICLs during DNA repair and may also have cross-link repair function that is independent of proteins in the Fanconi anemia (FA) pathway of DNA damage response and repair [5]. Importantly, FAN1 mutations are thought to lead to the renal disease Karyomegalic Interstitial Nephritis (KIN) [6-8].

The crystal structures of human FAN1 (hFAN1) and a bacterial FAN1 ortholog have been solved by several groups [9-11]. These structures provide insights into how FAN1 binds 5’ flap DNA with a 1-nt overhang (Fig. 1A). Surprisingly, the two available structures of the hFAN1 differ both in the oligomerization state (dimer vs monomer) and DNA-binding mode. The dimeric structure has an interface with DNA that spans both hFAN1 protomers [9], while the monomeric hFAN1 structure largely resembles the crystal structure of the *Pseudomonas aeruginosa* FAN1 (PaFAN1) bound to DNA [10,11] (Fig. 1A, right panel). It is unclear whether the two hFAN1 crystal structures capture physiologically relevant oligomeric states of this enzyme in different substrate engagement/cleavage modes.

**Fig. 1.**
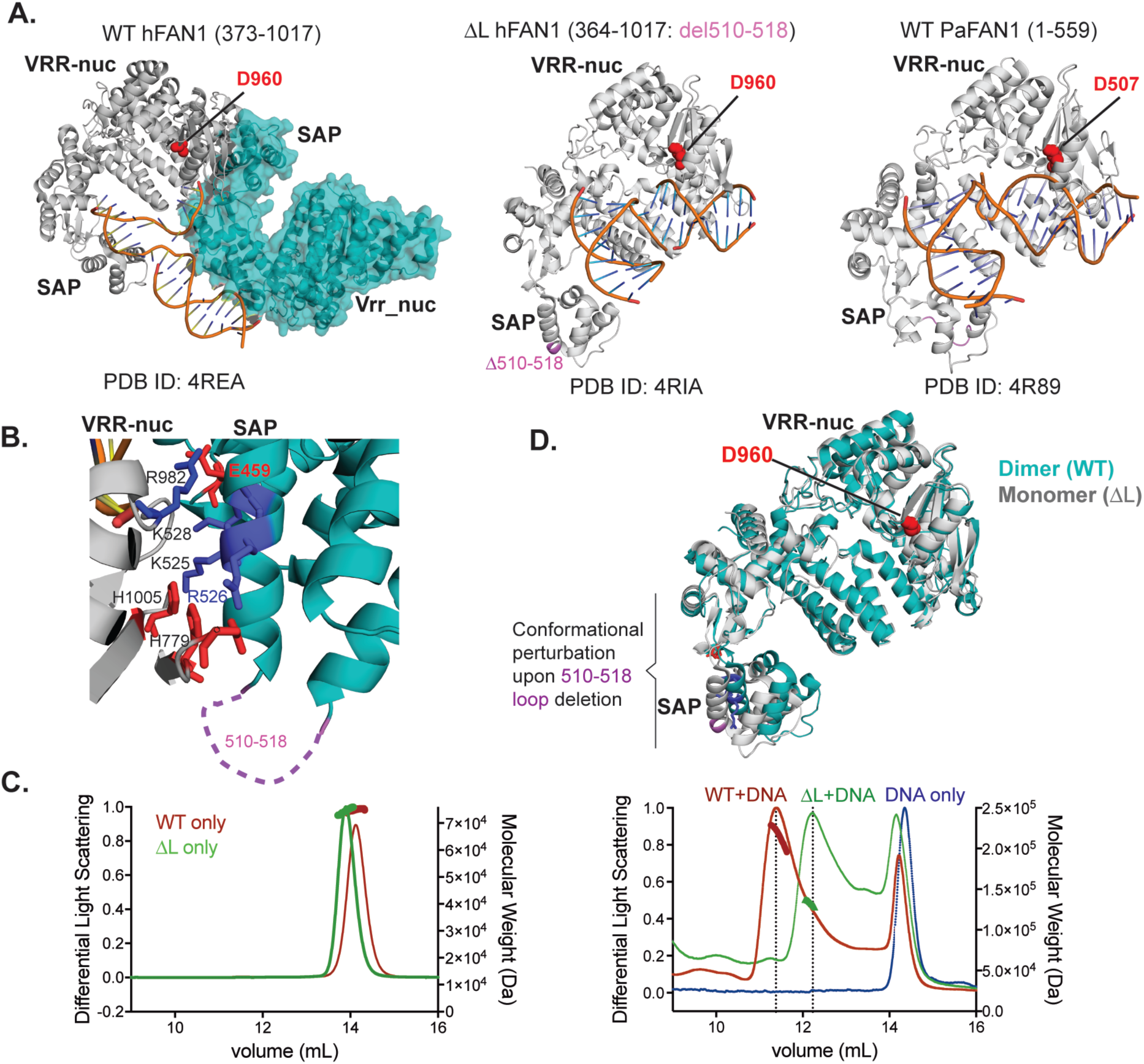
Dimerization of FAN1 upon DNA binding. (A) Previously solved crystal structures of WT hFAN1 [9], hFAN1 with residues 510-518 deleted ( Δ L hFAN1) [10], and *Pseudomonas aeruginosa* FAN1 [11] are compared, with the catalytic site aspartate highlighted in red. The catalytic VRR-nuc domain and the DNA binding SAP domain are labeled in bold for all FAN1 molecules. (B) Zoomed-in view of the dimerization interface of WT hFAN1, formed by the VRR-nuc domain of one monomer (grey) and the SAP domain of the 2^nd^ monomer (teal). The location of the 510-518 loop is marked with a dash line. (C) Multiangle laser light scattering (MALLS) chromatographic profiles for WT and ΔL hFAN1 alone (left panel) and with a 5’ flap substrate added (right panel). (D) Superposition of the monomeric FAN1 structure (PDB ID: 4RIA, grey) over one molecule of the dimeric FAN1 structure (PDB ID: 4REA, teal) highlighting the conformational perturbation induced by deletion of the flexible loop between residues 510-518.

In this study, we resolve the activity differences between the dimeric and monomeric forms of hFAN1 in the cleavage of 5’ DNA flaps. Interestingly, we find that even though both dimeric and monomeric hFAN1 forms can cleave a short DNA flap strand efficiently, monomeric hFAN1 is less capable than the dimeric enzyme in cleaving a longer flap. We independently verify this activity profile using bacterial orthologs of FAN1 that are intrinsically dimeric or monomeric.

## RESULTS AND DISCUSSION

### Requirement for an interdomain loop in DNA-induced hFAN1 dimerization

The crystal structure of hFAN1 previously solved by us shows a homodimeric configuration of the protein [9], where the two hFAN1 molecules make contacts with the duplex regions of the 5’ flap DNA so as to position one of the subunits to cleave the single-stranded DNA flap. This DNA-bound hFAN1 structure was derived from a truncated form of hFAN1 (residues 373-1017) that lacks the UBZ domain but otherwise covers all the essential domains for DNA processing and possesses nuclease activity comparable to that of full length hFAN1. We refer to this construct as WT hFAN1 or simply WT (Fig. 1A, left panel and Fig. 1B). Intriguingly, another published hFAN1 structure shows a monomeric conformation and a different mode of DNA binding [10]. This structure was similarly derived from hFAN1 (residues 364-1017) that lacks the UBZ domain but in addition harbors the deletion of a flexible loop spanning residues 510-518 in the SAP domain, hereafter referred to as ΔL hFAN1 or simply ΔL (Fig. 1A, middle panel).

To confirm that dimeric and monomeric hFAN1 forms exist in solution, we used size exclusion chromatography followed by multi-angle laser light scattering (SEC-MALLS) to characterize the WT and ΔL proteins. The data were collected either for the proteins alone or in the presence of a 5’ flap DNA substrate (Fig. 1C). We found that both WT and ΔL hFAN1 are monomeric in solution with a measured molecular weight of ~75 kDa. Upon pre-incubation with a 5’ flap DNA substrate, however, the SEC elution peak shifted to the dimeric position for WT, but the ΔL form remained monomeric. This provides direct evidence that deletion of the flexible loop prevents hFAN1 dimerization upon DNA binding.

We note that the results from the SEC-MALLS analysis are consistent with the proximal location of the flexible loop to the dimer interface in our hFAN1 structure (Fig. 1B). As a result, deletion of this loop (510-518) induces changes in the relative placement of the two helices abutting the loop (Supplemental Table 1) and a significant perturbation in the domain orientation of the hFAN1 molecule. Disruption of the dimer structure gives rise to the different DNA binding modes, especially at the SAP domain of the molecule (Fig. 1D), as observed in the two published X-ray crystal structures [9,10]. Based on the results above, we conclude that hFAN1 dimerization is dependent on conformational changes in the interface surrounding the flexible loop. Moreover, we hypothesize that the hFAN1 dimer represents an important physiological form of the enzyme.

### Flap nuclease activity profile of WT and ΔL hFAN1

Next, we tested the nuclease activity of WT and ΔL forms of hFAN1 on different 5’ flap substrates in time-course experiments. We found that the cleavage pattern for WT and ΔL is the same for flap substrates that harbor a 5’ overhang of either 1 nt or 5 nt (Fig. 2A, B), and nearly complete substrate cleavage occurs with both hFAN1 forms within 2 min. However, as the 5’ overhang length was increased to 15 nt and then to 40 nt (Fig. 2C, D), the cleavage efficiency diminished significantly for both WT and ΔL, albeit the activity drop on the 40-nt flap was much more pronounced for ΔL than for WT. These initial assays were performed using the UBZ domain-deleted (ΔUBZ) version of WT and ΔL hFAN1, as the overall purification yield of the ΔUBZ variant of these proteins was significantly higher than that of full-length hFAN1. To determine whether the UBZ domain exerts any influence on the flap nuclease activity of FAN1, we also expressed and purified full-length WT and ΔL FAN1 proteins and tested them on the same flap substrates (Fig. 2E-F). The results confirmed that while the ΔL mutant is just as capable as the WT counterpart in cleaving the 1-nt flap, the latter is much more active on the 40-nt flap than the mutant (Fig. 2E right panel).

**Fig. 2.**
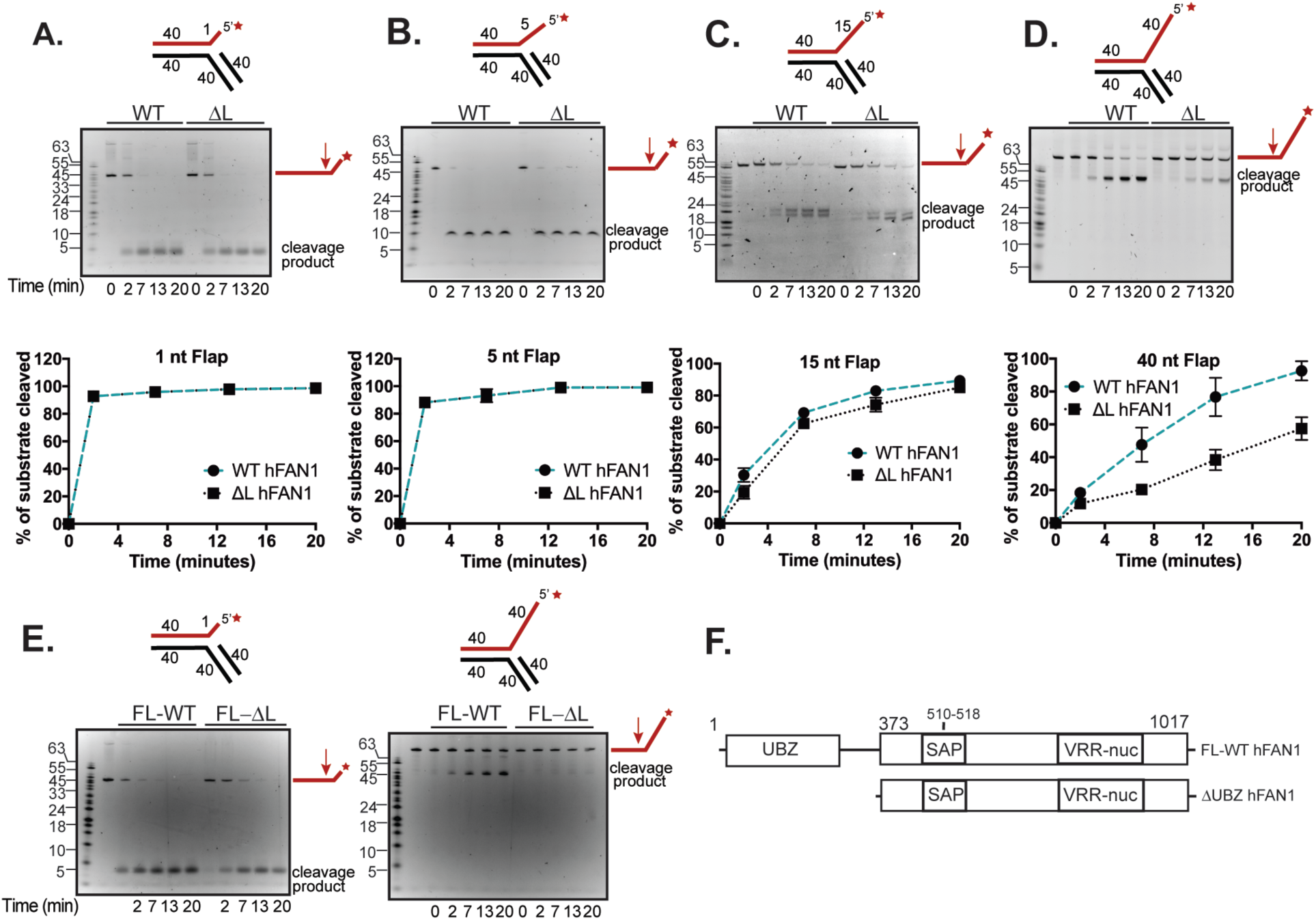
WT hFAN1 is more active on 5’ flap substrates with long overhang than ΔL hFAN1. In (A) to (D), the top panel shows the 5’ flap DNA substrate with dsDNA arms being fixed at 40 base pairs and the 5’ ssDNA overhang ranging from 1 nucleotide (nt) to 40 nt. The analysis of the time course reaction is shown in the middle panel. The fluorescently labeled strand is shown in “red” with the asterisk denoting the labeled end and the red arrow marking the cleavage site. The data were quantified and plotted in the bottom panel (n=3 for error bars; error bar = mean with SEM). (E) The nuclease activity of full-length hFAN1 and ΔL-hFAN1 was analyzed using DNA substrates with the 1-nt (left) or 40-nt (right) overhang. (F) Schematic of the hFAN1 proteins used in this study.

### Analysis of monomeric and dimeric bacterial FAN1 proteins for DNA flap cleavage

*Vibrio vulnificus* FAN1 (VvFAN1) is constitutively dimeric, while *Pseudomonas aeruginosa* FAN1 (PaFAN1) is monomeric. These enzymes have SAP and VRR-nuc domains similar to hFAN1, but both lack the ubiquitin-binding UBZ domain found in the human counterpart (Fig. 3A, left panel). Firstly, we used size exclusion chromatography to confirm that PaFAN1 is monomeric with and without DNA being present (Fig. 3A, right panel). On the other hand, we found that VvFAN1 is dimeric regardless of whether DNA is present or not (Fig. 3A, middle panel). Next, we tested these two bacterial proteins in time-course experiments with the same series of 5’ flap DNA substrates as what we used for hFAN1. The results revealed that PaFAN1 and VvFAN1 both have robust activity on substrates with 1-nt or 5-nt overhang, but increasing the length of the overhang to 15-nt or 40-nt leads to a reduction in DNA incision by both proteins, but with monomeric PaFAN1 being affected to a significantly higher degree (Fig. 3B-E). This trend recapitulates our results from examining ΔL hFAN1 and WT hFAN1.

**Fig. 3.**
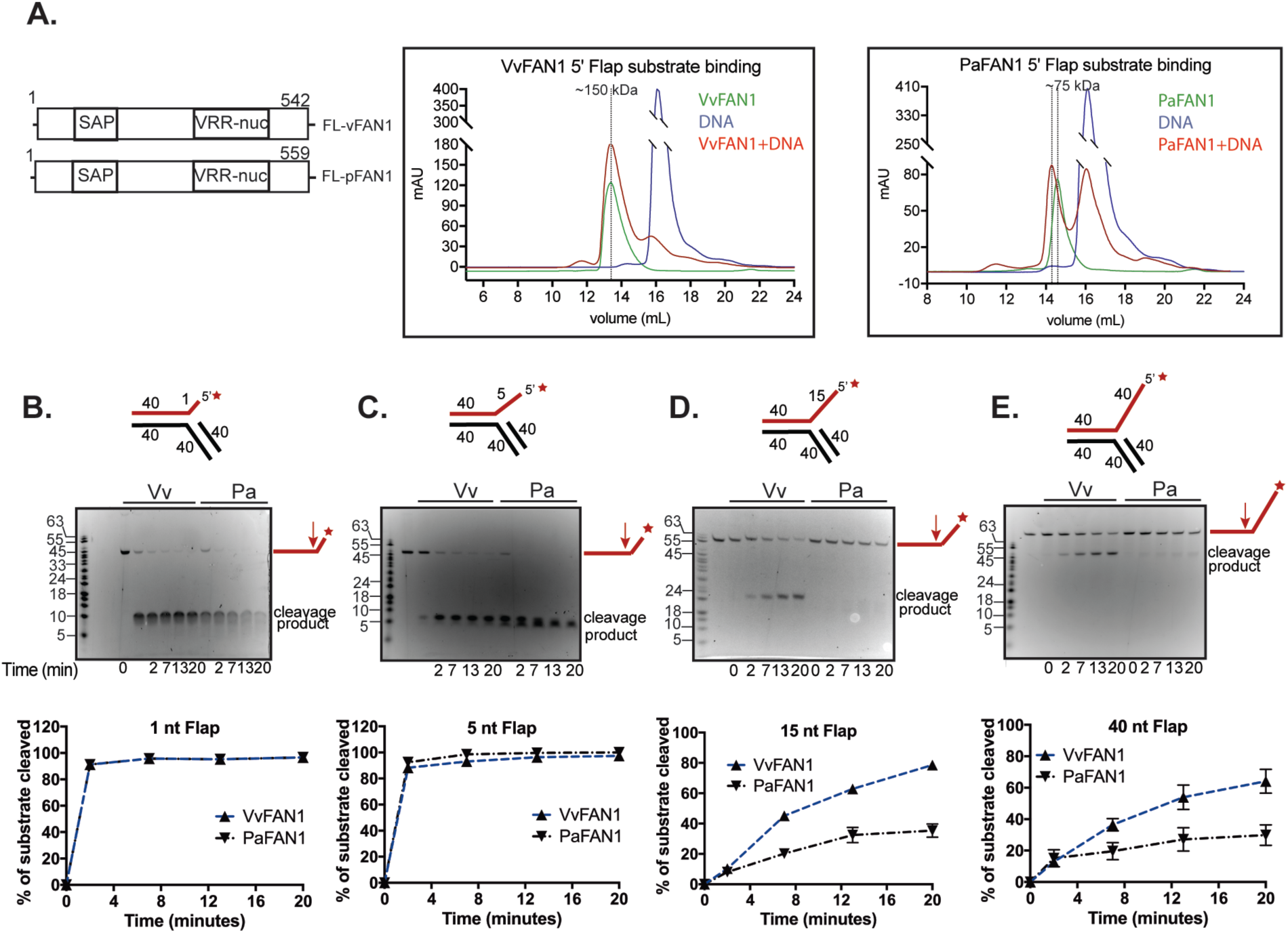
Analysis of bacterial dimeric and monomeric FAN1 proteins for 5’ DNA flap cleavage. (A) Schematic of the bacterial FAN1 proteins examined (left). Size exclusion chromatography profiles of VvFAN1 alone and bound to 5’ flap substrate (middle panel) and of PaFAN1 alone and bound to 5’ flap substrate (right panel). (B) to (E) The nuclease activity of VvFAN1 and PaFAN1 was tested in time course experiments using the flap substrates with 1-, 5-, 15- or 40-nt 5’ ssDNA overhang. See Figure 2 (A) to (D) for details.

## CONCLUSION

Important strides towards understanding FAN1’s function were made by solving the high-resolution X-ray crystal structures of human and bacterial FAN1s [9-11] (Fig. 1A). Intriguingly, these structures also raised the question about the relevant oligomeric state of FAN1 in human cells. In the present study, we provide data to clarify the major difference between dimeric and monomeric forms of FAN1 and to advance our understanding of the biological function of FAN1. Specifically, given the very different modes of DNA binding in the two hFAN1 structures, we predicted that a difference should manifest in DNA cleavage as well. Through testing a series of 5’ flap DNA substrates, we found that as the ssDNA overhang length increases, the efficiency of cleavage by FAN1 is reduced and that this reduction is significantly more dramatic for the monomeric form of hFAN1 (Fig. 4A). To ascribe this activity difference to the oligomeric status difference of the protein, we also examined bacterial FAN1 orthologs that are either dimeric or monomeric. Importantly, by comparing the dimeric VvFAN1 and the monomeric PaFAN1 in the same nuclease tests, we have provided further evidence that the oligomeric status of FAN1 determines its efficiency in cleaving a long ssDNA flap strand.

**Fig. 4.**
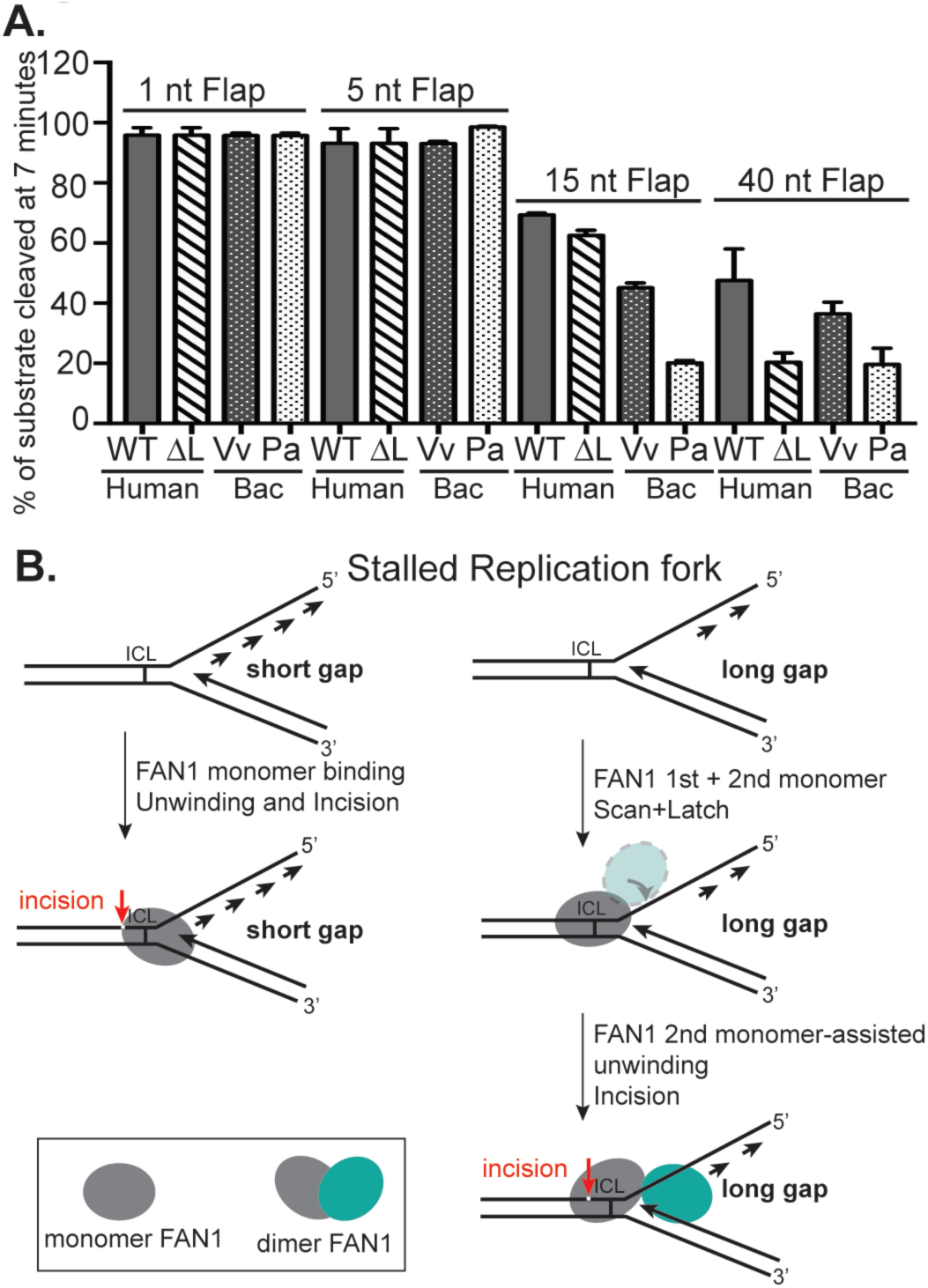
Dimeric FAN1 is the relevant form for cleavage of long DNA flaps. (A) Summary of cleavage of substrates with different overhang lengths by WT and ΔL hFAN1 and by bacterial FAN1s [Vibrio (Vv) and Pseudomonas (Pa)] at 7-minute time point. (B) Model illustrating how protein dimerization activates ICL incision at stalled replication fork with a long ssDNA gap.

Based on our results, we surmise that cleavage past an ICL that abuts a short ssDNA gap at a replication fork can be carried out efficiently by either dimeric or monomeric hFAN1, whereas the equivalent reaction involving replication forks with a larger DNA gap is likely catalyzed by the dimeric form of the enzyme (Fig. 4B). Our crystal structures of dimeric hFAN1 have provided evidence for “scanning”, “latching” and “unwinding” modes of DNA engagement stemming from the concerted effort of two FAN1 protomers [9](Fig. 4B). We note that the VRR-nuc domains form a dimeric structure that can cleave the Holliday Junction in a symmetric fashion, but due to the insertion of a helix in the VRR-nuc domain of FAN1, the inherent dimerization property becomes impaired [12]. In this regard, certain FAN1 orthologs, particularly the human and Vibrio enzymes, have evolved another surface for protein dimerization where the SAP domain of one protomer associates with the VRR-nuc domain of a second protomer to generate an asymmetric dimer. This asymmetry underlies the ability of hFAN1 to cleave a long 5’ flap strand. Thus, protein dimerization likely imparts plasticity to hFAN1 in the processing of damaged replication forks.

## EXPERIMENTAL PROCEDURES

### Protein expression vectors

The expression vectors for human FAN1 (hFAN1), the UBZ domain deleted hFAN1_373-1017_ and the ΔL variants have been described [2,9]. The *Pseudomonas aeruginosa* FAN1 (PaFAN1) and *Vibrio vulnificus* FAN1 (vFAN1) coding sequences were inserted into the pET28a vector, with 6XHis-SUMO coding sequence fused to the N-terminus of both sequences.

### Protein purification

The hFAN1 species were purified using the protocol reported previously [2]. hFAN1_373-1017_ and its ΔL variant were expressed in BL21 (DE3) *E. coli* cells by pre-induction growth at 37°C until O.D._600_ of 0.8, followed by protein induction with 0.2 mM isopropyl b-D-thiogalactopyranoside (IPTG) at 16°C for 16 h. Cells were harvested by centrifugation, resuspended in 1X PBS buffer with 1M sodium chloride, 5% Glycerol and 1 mM DTT, followed by sonication to lyse the cells. After removing cell debris by centrifugation at 25,000 xg for 1 h, the supernatant was subjected to affinity purification with Nickel-NTA beads (GE Healthcare). On-column cleavage of His-MBP fusion protein was done by overnight incubation with SARS-CoV M-pro protease [13], followed by elution, size exclusion chromatography purification, and finally pooling of FAN1 fractions and concentrated to 5 mg/ml before freezing (REFs). PaFAN1 and Vv FAN1 were expressed similarly as human FAN1 proteins. Cells were harvested by centrifugation, resuspended in 1X PBS buffer with 1M sodium chloride, 5% Glycerol and 1 mM DTT, followed by sonication to lyse the cells. After removing cell debris by centrifugation at 25,000 xg for 1 h, the supernatant was subjected to affinity purification with Nickel-NTA beads. PaFAN1 and VvFAN1 were eluted off the Ni-NTA with imidazole and subjected to overnight His-SUMO cleavage by the Ulp1 protease. The cleavage mix was subjected to size exclusion chromatography, followed by pooling of the FAN1 peaks, concentration to 5 mg/ ml and storage at -80°C.

### DNA substrates

All the oligonucleotides (Supplemental Table 2) were purchased from Integrated DNA Technologies Inc. Selected oligonucleotides were purchased pre-labeled with 6-FAM (Fluorescein) at either the 5’ or 3’ end. The 5’ flap substrates (Supplemental Table 3) were made by heating a mixture of the three constituent oligonucleotides (final concentration of 10 μM each) in 20 mM Tris-HCl pH 7.5, 50 mM NaCl for 5 min at 95°C, followed by slow cooling to room temperature. The substrates were stored at -20°C.

### Multi-angle laser light scattering (MALLS) analysis

MALLS was performed in line with a Superdex 200 10/300 size exclusion chromatography column (GE Healthcare) on a Ettan LC system (GE Healthcare) in buffer A (1X phosphate buffered saline, 1M NaCl and 0.1 mM TCEP) for FAN1 without DNA and in buffer B (1X phosphate buffered saline, 50 mM NaCl, 1 mM EDTA and 0.1 mM TCEP) for FAN1 with 10,10,1dT DNA (Supplemental Tables 2 & 3). The system was coupled on-line to an 18-angle MALLS detector and a differential refractometer (DAWN HELEOS II and Optilab rEX, Wyatt Technology). Molar mass determination was calculated with the ASTRA 6.2 software.

### Nuclease assay

The nuclease activity of human and bacterial FAN1 proteins was examined according to published procedures [5,11]. Reaction mixtures were resolved in 15% denaturing polyacrylamide gels (with 8 M Urea) in 1X TBE buffer (89 mM Tris Borate, pH 8.4, and 2 mM EDTA) at 200 V for 40 min at room temperature. The fluorescently labeled DNA species were visualized in a G:box (Syngene). Band intensities were quantitated with the ImageJ software [14].

## Abbreviations

FAN1: FANCD2/FANCI-associated nuclease 1
FANCI: Fanconi anemia complementation group I
FANCD2: Fanconi anemia complementation group D2
ICL: Interstrand DNA crosslink
HR: homologous recombination
TLS: translesion DNA synthesis

## SUPPLEMENTAL INFORMATION

Supplemental file includes 3 tables

## ACKNOWLEDGMENTS

We thank William Eliason for technical support in the SEC-MALLS analysis. This work was supported by National Institutes of Health Grants R01 ES007061, RO1 CA168635, R35 GM118047 and a Young Investigator Award from the Alex’s Lemonade Stand Foundation for Childhood Cancer.

## CONFLICT OF INTEREST

The authors declare that they have no conflict of interest with the contents of this article. The content is solely the responsibility of the authors and does not necessarily represent the official views of the National Institutes of Health.

## AUTHOR CONTRIBUTIONS

T.R. and S.L. designed the experiments and T.R did protein expression, purification, DNA binding, nuclease activity assays. W.Z. and H.A. helped with data analysis. T.R. wrote the manuscript with input from Y.X. and P.S.

